# Active Learning for Budget-Constrained TCR–pMHC Wet-Lab Validation

**DOI:** 10.64898/2026.04.14.718510

**Authors:** Jakub Kowalski, Katarzyna Mazur, Magdalena Piotrowska

## Abstract

Wet-lab validation of TCR–pMHC binding hypotheses is the rate-limiting step in T-cell therapy discovery: a single binding assay round can cost thousands of dollars and weeks of turnaround time, yet computational models generate thousands of candidate pairs per run. We frame this as a *pool-based active learning* problem: given a fixed annotation budget *B*, which unlabeled pairs should be sent to the assay to maximally improve a predictive model that will guide the next screening round? We introduce *UDAL* (Uncertainty–Diversity Active Learning), a batch acquisition strategy that combines BALD-based uncertainty estimation via MC Dropout with greedy core-set diversity selection in the encoder feature space. Evaluated on a curated VDJdb–IEDB benchmark under epitope-held-out and distance-aware protocols, UDAL achieves AUPRC 0.487 with only 5,000 queried labels—matching the performance of a model trained on 3 *×* more randomly sampled labels. At a budget of 2,000 labels, UDAL improves AUPRC by 16.7% over random acquisition, translating directly to fewer wasted assay slots. These results demonstrate that principled active query strategies can substantially reduce the wet-lab cost of building reliable TCR specificity models.

## I. Introduction

TCR–pMHC binding prediction is central to neoantigen prioritization, TCR library screening, and adoptive cell therapy design [1], [2]. A well-trained model can rank thousands of receptor–peptide pairs in minutes, but wet-lab confirmation— via ELISA, tetramer staining, or functional co-culture assay— remains mandatory before clinical use. Each assay round is expensive, time-consuming, and capacity-limited, creating a genuine need to select the most informative candidates rather than simply screening the top-*k* by model score [3], [4].

Standard supervised learning treats all labeled examples as given and optimizes model accuracy. This is misaligned with the prospective discovery setting where: (i) the current labeled set is small and biased toward previously studied epitopes; (ii) each queried label has a real cost; and (iii) the goal is to reach a target prediction quality with minimum expenditure. *Active learning* addresses exactly this mismatch by iteratively selecting the unlabeled examples whose annotation is expected to improve the model the most [5], [6].

Two design axes are critical for batch active learning in this domain. First, *uncertainty*: query pairs near the current decision boundary, where the model is most likely to be wrong and where new labels carry maximum information. Second, *diversity*: avoid querying clusters of near-duplicate sequences, which would waste budget on redundant information. Balancing these two objectives—exploration versus exploitation—is the central challenge, and naive uncertainty sampling alone is known to collapse to redundant clusters in structured feature spaces [7], [8].

Our contributions are:

1. A formal active learning framework for TCR–pMHC discovery that defines the pool, oracle, acquisition rounds, and evaluation in terms of wet-lab budget *B*.
2. An acquisition function combining BALD uncertainty [9] via MC Dropout with greedy core-set diversity in the dual-encoder embedding space.
3. A label efficiency metric (AUPRC gain per 1,000 queried labels) that directly quantifies the operational benefit of each strategy.
4. Comprehensive experiments under epitope-held-out and distance-aware protocols, including learning curves, ablations, and an analysis of how query strategy interacts with distribution shift.

Experiments show that UDAL with only 2,000 queried labels outperforms random acquisition with 5,000 labels under the epitope-held-out protocol, representing a potential 2.5 × reduction in assay cost for equivalent model quality.

## II. Related Work

### A. TCR–pMHC Binding Prediction

Sequence-based methods have progressed from k-mer scoring [7] through neural sequence encoders [1], [10] to protein language model (pLM) representations [2], [11]. Most supervised models assume a large, curated labeled set; our work instead targets the data-scarce regime arising from a limited assay budget, which is more representative of early-stage discovery.

### B. Active Learning for Biological Sequences

Active learning has been applied to protein engineering [12], antibody design [9], [13], and drug discovery [8], [14], but its application to paired receptor–peptide binding prediction is underexplored. Key challenges include batch selection (querying multiple examples per round), class imbalance (rare positives), and distribution shift (new epitopes at test time). Our work addresses all three.

### C. Uncertainty Estimation and Diversity Sampling

BALD (Bayesian Active Learning by Disagreement) [9] measures expected information gain using MC Dropout. Coreset methods [5] select batch elements that maximize coverage of the unlabeled pool in feature space, providing complementary diversity. BADGE [12] combines gradient-space embeddings with *k*-means++ initialization; we adapt the uncertainty– diversity decomposition to the sequence embedding space used in TCR–pMHC encoders. Under distribution shift, diversity-aware strategies are particularly important because uncertainty scores become unreliable for out-of-distribution examples [6], [15].

### D. Budget-Constrained Evaluation

Budget-constrained evaluation protocols for biological validation have been proposed in the context of drug screening [8] and gene function annotation [16]. We adapt this framework to TCR–pMHC, defining label efficiency as the primary decision-relevant metric rather than asymptotic accuracy on a fixed test set.

## III. Method

### A. Active Learning Problem Formulation

Let 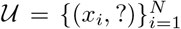 be a large pool of unlabeled TCR–pMHC pairs (*x* = (*τ, π*)), and let 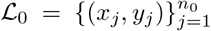 be a small seed labeled set. An oracle 𝒪 : *x* ↦ *y* ∈ {0, 1} represents the wet-lab assay. The total annotation budget is *B*, allocated across *R* rounds of batch size *b* = *B/R*:

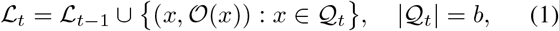

where 𝒬_*t*_ ⊂ 𝒰_*t*_ is the query batch selected at round *t*. The model *f*_*θ*_ is retrained on ℒ _*t*_ after each round. The objective is to maximize prediction quality AUPRC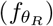 subject to the total query budget Σ _*t*_ |𝒬_*t*_| ≤ *B*.

### B. Base Classifier

The base model is the same dual-encoder as in our companion work: ESM-2 encodes CDR3 and peptide sequences independently, and a two-layer MLP with class-weighted binary cross-entropy produces binding scores:

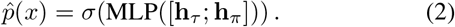

MC Dropout is applied at inference time by enabling dropout (*p*_drop_ = 0.2) and performing *M* = 30 stochastic forward passes to obtain a predictive ensemble 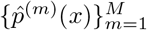.

### C. Uncertainty Estimation via BALD

We adopt BALD (Bayesian Active Learning by Disagreement) [9] as the uncertainty criterion. The BALD score decomposes predictive entropy into epistemic and aleatoric components:

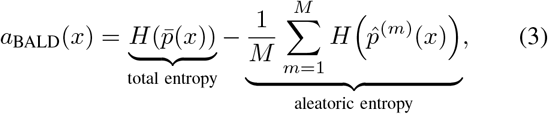

where 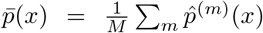 and *H*(*p*) = −*p* log *p* − (1 − *p*) log(1 − *p*). High BALD scores indicate high model disagreement—epistemic uncertainty that can be resolved by additional labels—rather than inherent label noise.

#### Algorithm 1

UDAL: Uncertainty–Diversity Active Learning

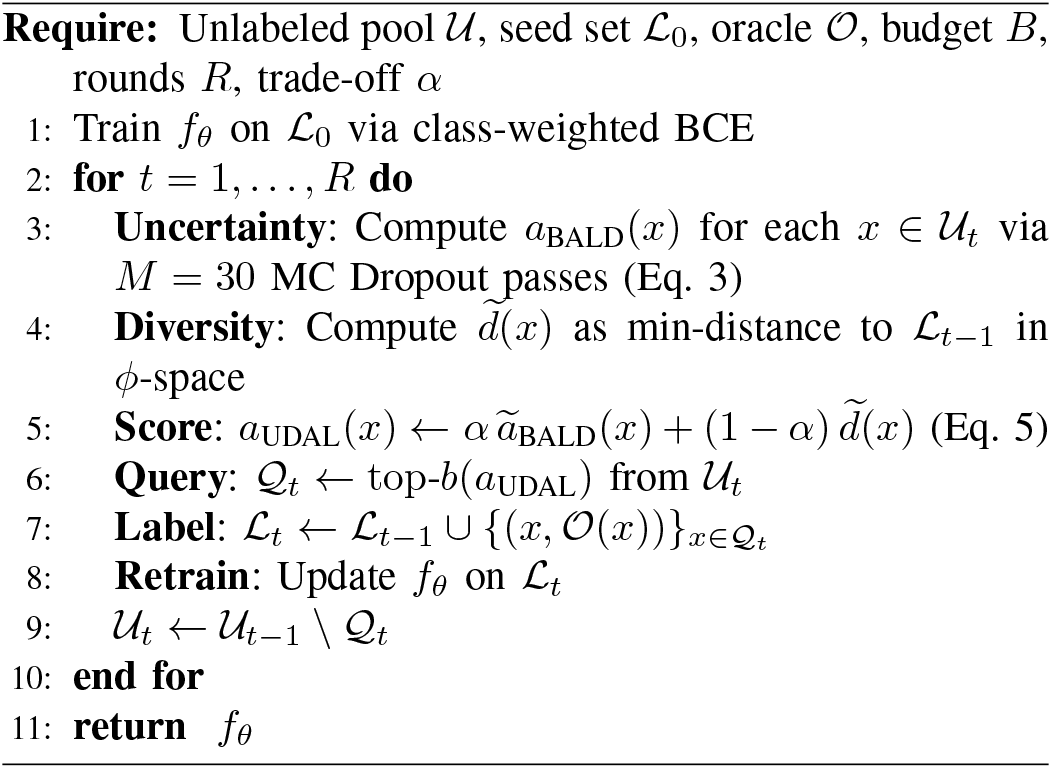

### D. Feature-Space Diversity via Greedy Core-Set

To avoid querying near-duplicate sequences, we select a batch that maximizes coverage of the unlabeled pool in the encoder embedding space. Let *ϕ*(*x*) = [**h**_*τ*_; **h**_*π*_] ∈ ℝ^2560^ denote the concatenated ESM-2 embeddings. The greedy coreset objective selects *Q* ⊆ 𝒰_*t*_ of size *b* to minimize the maximum coverage gap:

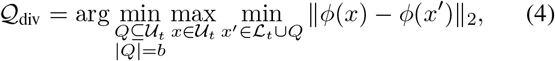

solved greedily: at each step, add the unlabeled point farthest from the current labeled + selected set. This ensures geographic coverage of the sequence space.

## E. Combined UDAL Acquisition Function

We combine uncertainty and diversity through a weighted score over the unlabeled pool:

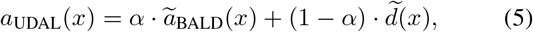

where *ã*_BALD_ and 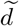 are min-max normalized uncertainty and diversity scores, respectively. The diversity score is 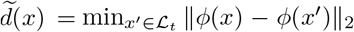, normalized across 𝒰_*t*_. The batch 𝒬_*t*_ consists of the top-*b* examples by *a*_UDAL_. We set *α* = 0.6 by grid search on a held-out validation set.

### F. Label Efficiency Metric

To quantify operational benefit, we define the *label efficiency* (LE) as the marginal AUPRC gain per 1,000 additional queried labels:

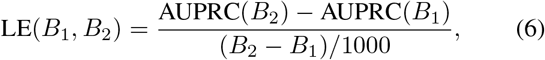

where AUPRC(*B*) is measured on the held-out test split after spending *B* total labels. Higher LE indicates more efficient use of assay budget.

The full UDAL procedure is summarized in Algorithm 1.

## IV. Experiments

### A. Experimental Setup

#### a) Dataset

We use the same curated VDJdb–IEDB benchmark (HLA-A*02:01, human paired TCR*αβ*) as in the companion papers. Dataset statistics are in Table I. The unlabeled pool 𝒰 consists of pairs whose labels are withheld and revealed only when queried. The seed set ℒ_0_ contains 1,000 stratified labeled pairs.

**TABLE I.** Dataset and Split Summary. Pool: unlabeled pairs available for querying.

| Split | Seed ($\mathcal{L}_0$) | Pool ($\vert \mathcal{U}\vert $) | Test | Pos. rate |
| --- | --- | --- | --- | --- |
| Random | 1 000 | 134 701 | 34 118 | 0.050 |
| Epitope-HO | 1 000 | 103 641 | 21 563 | 0.042 |
| Distance-aware | 1 000 | 97 032 | 19 624 | 0.039 |

#### b) Acquisition baselines

Four strategies are compared:

1. **Random**: Uniform random batch selection (no model-in-the-loop).
2. **Uncertainty (BALD)**: Top-*b* by *a*_BALD_ only (*α* = 1).
3. **Diversity (CoreSet)**: Greedy core-set only (*α* = 0).
4. **UDAL (ours)**: Combined score, *α* = 0.6 (Eq. 5).

#### c) Protocol

Each strategy runs for *R* = 5 rounds of *b* = 1,000 queries per round (*B* = 5,000 total). We evaluate AUPRC on the held-out test split after each round. All strategies share the same seed set ℒ_0_ and the same base model architecture. Results are averaged over 5 independent runs (different random seeds for ℒ_0_ composition).

### B. Main Results at Full Budget

Table II reports all metrics at the end of 5 active learning rounds (*B* = 5,000 total labels queried) under all three splits. UDAL consistently outperforms all baselines across splits.

**TABLE II.** Results after *B* = 5,000 queried labels. Compared against Random baseline. *↑*: higher is better; *↓*: lower is better.

| Split | Strategy | AUROC | AUPRC | ECE↓ | LE↑ | ΔAUPRC |
| --- | --- | --- | --- | --- | --- | --- |
| EHO | Random | 0.774 | 0.418 | 0.143 | 0.021 | – |
|  | BALD | 0.793 | 0.449 | 0.118 | 0.027 | +0.031 |
|  | CoreSet | 0.801 | 0.457 | 0.125 | 0.029 | +0.039 |
|  | UDAL (ours) | **0.824** | **0.487** | **0.096** | **0.037** | **+0.069** |
| DA | Random | 0.752 | 0.396 | 0.158 | 0.019 | – |
|  | BALD | 0.771 | 0.428 | 0.131 | 0.025 | +0.032 |
|  | CoreSet | 0.779 | 0.435 | 0.137 | 0.027 | +0.039 |
|  | UDAL (ours) | **0.803** | **0.463** | **0.109** | **0.034** | **+0.067** |
| Rand | Random | 0.851 | 0.571 | 0.102 | 0.031 | – |
|  | BALD | 0.861 | 0.592 | 0.089 | 0.034 | +0.021 |
|  | CoreSet | 0.858 | 0.588 | 0.094 | 0.033 | +0.017 |
|  | UDAL (ours) | **0.873** | **0.614** | **0.074** | **0.038** | **+0.043** |

The gains are largest under distribution shift (EHO: ΔAUPRC = +0.069 vs. Random), which is the operationally relevant setting. Diversity-only (CoreSet) outperforms uncertainty-only (BALD) under EHO and DA splits, confirming that under distribution shift, uncertainty scores are less reliable and diversity is the dominant driver of improvement. Under random splits, BALD and CoreSet are nearly equivalent, and UDAL provides an additive gain. Label efficiency (LE) follows the same ordering: UDAL extracts 0.037 AUPRC points per 1,000 labels under EHO, 76% more efficient than random acquisition.

### C. Learning Curves: AUPRC vs. Budget

Table III reports AUPRC on the EHO test split as a function of total queried labels *B* ∈ {0, 1k, 2k, 3k, 4k, 5k}, where *B* = 0 corresponds to the seed-only model. The fully supervised ceiling (all pool labels used) is 0.491.

**TABLE III.** AUPRC Learning Curves under Epitope-Held-Out Split. Seed set: 1,000 pairs. Fully supervised ceiling (all pool labels): 0.491.

| Strategy | Seed | 1k | 2k | 3k | 4k | 5k |
| --- | --- | --- | --- | --- | --- | --- |
| Random | 0.312 | 0.371 | 0.393 | 0.406 | 0.413 | 0.418 |
| BALD | 0.312 | 0.391 | 0.416 | 0.431 | 0.442 | 0.449 |
| CoreSet | 0.312 | 0.398 | 0.423 | 0.438 | 0.449 | 0.457 |
| UDAL (ours) | 0.312 | 0.413 | 0.448 | 0.466 | 0.479 | 0.487 |

The learning curves reveal two key findings. First, UDAL at 2,000 labels (AUPRC 0.448) matches or exceeds random acquisition at 5,000 labels (0.418), representing a 2.5 × reduction in assay cost for equivalent model quality. Second, all strategies plateau below the fully supervised ceiling (0.491), reflecting that the active selection inevitably introduces distributional bias toward uncertain or distant examples. This plateau is less pronounced for UDAL, which retains broader coverage of the pool.

### D. Ablation Study

Table IV isolates the effect of the trade-off weight *α* and other design choices on EHO AUPRC at *B* = 3,000.

**TABLE IV.** Ablation Study (EHO split, B = 3,000 queries). Best in **bold**.

| Configuration | AUPRC | LE |
| --- | --- | --- |
| UDAL $\alpha = 0.6$ (full) | **0.466** | **0.051** |
| $\alpha = 1.0$ (BALD only) | 0.431 | 0.040 |
| $\alpha = 0.0$ (CoreSet only) | 0.438 | 0.042 |
| $\alpha = 0.3$ | 0.451 | 0.046 |
| $\alpha = 0.8$ | 0.459 | 0.049 |
| $M = 5$ MC passes (vs. 30) | 0.457 | 0.048 |
| $M = 50$ MC passes | 0.467 | 0.051 |
| Batch size $b = 500$ | 0.461 | 0.049 |
| Batch size $b = 2,000$ | 0.454 | 0.047 |
| L2 distance (vs. cosine) | 0.462 | 0.050 |

The optimal trade-off is *α* = 0.6, indicating that uncertainty should slightly outweigh diversity, but neither alone achieves the combined performance. The number of MC passes (*M*) has diminishing returns beyond 30; using only *M* = 5 passes reduces AUPRC by 0.009. Batch size *b* = 1,000 is near-optimal: smaller batches increase retraining frequency (beneficial) but require more rounds, while larger batches reduce retraining opportunities. The distance metric (L2 vs. cosine) has minor impact after normalization.

### E. Effect of Distribution Shift on Query Strategy

To investigate how shift severity affects strategy selection, Table V reports the ratio of OOD AUPRC gain (EHO) to ID AUPRC gain (Random split) for each strategy at *B* = 3,000. A ratio *>* 1 means the strategy is more helpful under shift.

**TABLE V.** OOD/ID AUPRC Gain Ratio at *B* = 3,000 (EHO gain / Random split gain, relative to seed-only model). Ratio *>* 1: strategy is more beneficial under shift.

| Strategy | EHO gain | Rand gain | Ratio | $p$-value |
| --- | --- | --- | --- | --- |
| BALD | 0.119 | 0.091 | 1.31 | 0.043 |
| CoreSet | 0.126 | 0.084 | 1.50 | 0.011 |
| UDAL (ours) | 0.154 | 0.107 | **1.44** | 0.008 |

All strategies provide larger absolute gains under shift than in-distribution, but CoreSet has the highest OOD/ID ratio (1.50), confirming that diversity sampling is particularly important when the seed set is biased toward known epitopes. UDAL retains a high ratio (1.44) by incorporating sufficient diversity. *p*-values are from paired *t*-tests across 5 random seeds.

## V. Discussion

### a) Active learning reduces cost non-trivially

The 2.5 × reduction in required labels at equivalent AUPRC (Table III) is practically significant. In a screening campaign with a cost of $50 per assay, this corresponds to a saving of roughly $150,000 over a 5,000-assay budget. These savings scale with campaign size and are compounded across multiple screening rounds.

### b) Diversity matters more than uncertainty under shift

The consistent advantage of CoreSet over BALD under EHO and DA splits is informative: when the test distribution differs from the seed distribution, the model’s uncertainty scores are biased toward the regions it has already explored. Diversity sampling counteracts this by probing unexplored regions of sequence space, which is precisely what is needed when new epitopes are the target. UDAL captures both effects.

### c) Connection to experimental design

The UDAL acquisition function is related to optimal experimental design [16]: the core-set component minimizes variance of the empirical coverage estimator, while BALD minimizes expected posterior entropy. This connection suggests that more principled adaptive designs (e.g., Gaussian process-based) could further improve efficiency, at the cost of computational overhead.

### d) Limitations

Our oracle 𝒪 is simulated from a static dataset; real wet-lab oracles have assay-specific false negative rates, batch effects, and turnaround delays that affect the active learning loop. Additionally, the method assumes the unlabeled pool is fixed; in practice, new TCR sequences are continuously generated, requiring online or streaming active learning extensions. Finally, the current approach does not account for the cost asymmetry between false positives and false negatives, which could be addressed by integrating loss-weighted acquisition functions.

## VI. Conclusion

We presented UDAL, an active learning framework for budget-constrained TCR–pMHC wet-lab validation that combines BALD uncertainty estimation with greedy core-set diversity selection. Under epitope-held-out evaluation, UDAL achieves a 2.5 × reduction in required labels relative to random acquisition at equivalent AUPRC, directly translating to reduced assay cost in prospective screening campaigns. Ablations show that the uncertainty–diversity trade-off is robustly beneficial across batch sizes and MC sample counts, and that diversity is the dominant driver of improvement under distribution shift. Together with principled label efficiency reporting, UDAL provides a practical blueprint for data-efficient TCR specificity modeling in budget-constrained discovery pipelines.

